# N-methyl-bacillithiol, a new metabolite discovered in the *Chlorobiaceae*, indicates that bacillithiol and derivatives are widely phylogenetically distributed

**DOI:** 10.1101/173617

**Authors:** Jennifer Hiras, Sunil V. Sharma, Vidhyavathi Raman, Ryan A. J. Tinson, Miriam Arbach, Dominic F. Rodrigues, Javiera Norambuena, Chris J. Hamilton, Thomas E. Hanson

## Abstract

Low-molecular weight (LMW) thiols are metabolites that mediate redox homeostasis and the detoxification of chemical stressors in cells. LMW thiols are also thought to play a central role in sulfur oxidation pathways in phototrophic bacteria, including the *Chlorobiaceae*. Fluorescent thiol labeling of metabolite extracts coupled with HPLC showed that *Chlorobaculum tepidum* contained a novel LMW thiol with a mass of 412 ± 1 Da corresponding to a molecular formula of C_14_H_24_N_2_O_10_S. These data suggested the new thiol is closely related to bacillithiol (BSH), the major LMW thiol from low G+C% Gram-positive bacteria. By comparing the as-isolated bimane adduct with chemically synthesized candidate structures, the *Cba. tepidum* thiol structure was identified as N-methyl-bacillithiol (N-Me-BSH), methylated on the cysteine nitrogen, a rarely observed modification in metabolism. Orthologs of bacillithiol biosynthetic genes in the *Cba. tepidum* genome were required for the biosynthesis of N-Me-BSH. Furthermore, the CT1040 gene product was genetically identified as the BSH N-methyltransferase. N-Me-BSH was found in all *Chlorobi* examined as well as *Polaribacter* sp. strain MED152, a member of the *Bacteroidetes.* A comparative genomic analysis indicated that BSH/N-Me-BSH is synthesized not only by members of the *Chlorobi*, *Bacteroidetes*, *Deinococcus*-*Thermus*, and *Firmicutes*, but also by *Acidobacteria*, *Chlamydiae*, *Gemmatimonadetes*, and *Proteobacteria.*

**Significance Statement:** Here, N-Me-BSH is shown to be a redox-responsive LMW thiol cofactor in *Cba. tepidum* and the gene, *nmbA*, encoding the BSH N-methyltransferase responsible for its synthesis is identified. The co-occurrence of orthologs to BSH biosynthesis genes and bacillithiol N-methyltransferase was confirmed to correctly predict LMW thiol biosynthesis in phylogenetically distant genomes. The analysis indicates that BSH/N-Me-BSH are likely the most widely distributed class of LMW thiols in biology. This finding sheds light on the evolution of LMW thiol metabolism, which is central to redox homeostasis, regulation and stress resistance in all cellular life. It also sheds light on a rare chemical modification. N-Me-BSH is the fourth instance of cysteinyl nitrogen methylation in metabolism. Identification of the BSH N-methyltransferase reported here will enable detailed *in vivo* and *in vitro* dissection of the functional consequences of this modification. As a standalone N-methyltransferase, NmbA may be useful as a component of constructed biosynthetic pathways for novel product (bio)synthesis.

## INTRODUCTION

Structurally diverse low molecular weight (LMW) thiols, compounds containing free sulfhydryl groups and less than 1000 Da in mass, are found in all nearly all cells examined where they maintain redox homeostasis and detoxify chemical stressors (1). Glutathione (L-γ-glutamyl-L-cysteinyl-glycine, GSH), the best studied LMW thiol (Fig. 1), is distributed widely in eukaryotes. However, not all prokaryotes produce and utilize GSH; it is restricted to the *Cyanobacteria* and certain *Proteobacteria* (1). In recent decades a range of alternative cysteine-containing redox balancing LMW thiols have been discovered in prokaryotes including mycothiol (MSH, (2)) and bacillithiol (BSH) (3).

**Figure 1.**
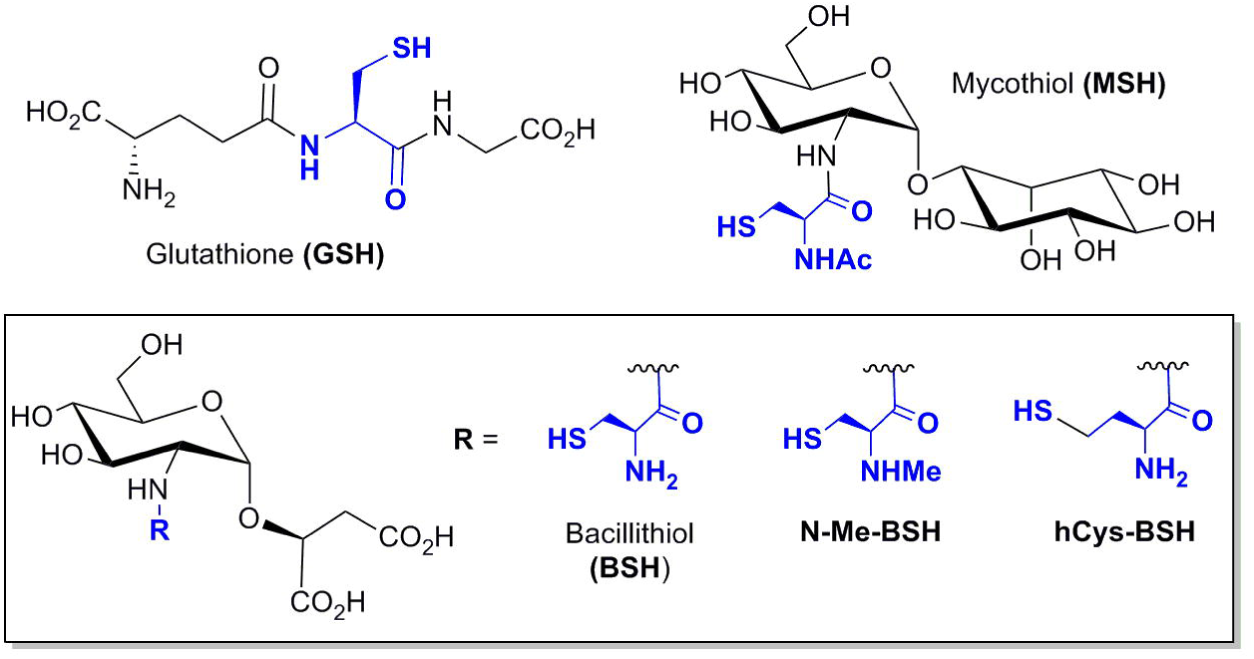
Structures of LMW thiols referred to in the text with the sulfhydryl bearing motif derived from cysteine or homocysteine colored blue. BSH derivatives are boxed together with the R group on the common backbone indicated.

The green sulfur bacteria (the *Chlorobiaceae*) are anaerobic, anoxygenic phototrophs that are found in anoxic water and sediments containing reduced sulfur compounds that are exposed to light. The *Chlorobiaceae* have contributed to our understanding of CO_2_ fixation via the reductive TCA cycle (4) and light-harvesting mechanisms through studies of the chlorosome (5, 6), the light harvesting antenna in this group. *Chlorobaculum tepidum* (formerly *Chlorobium tepidum*) is a model system for the *Chlorobi* because of its rapid growth rate (7), complete genome sequence (8), and genetic system (9–12).

*Cba. tepidum* oxidizes reduced sulfur compounds (sulfide, elemental sulfur and thiosulfate) to feed electrons into the photosynthetic electron transport chain where they ultimately reduce ferredoxin, which in turn is used to drive CO_2_ fixation (4), N_2_ fixation (13) and the reduction of NAD(P)^+^ (14). LMW thiols have been proposed as a sulfur atom shuttle between the periplasm and cytoplasm to feed sulfide into biosynthetic pathways, the dissimilatory sulfite reductase and ATP-sulfurylase in the *Chlorobiaceae* and other phototrophic sulfur oxidizers (15, 16). This function requires that the LMW thiol cycle between the thiol (R-SH) and perthiol (R-S_n_-SH) forms as observed for glutathione amide in the purple sulfur bacterium *Chromatium gracile* (17). However, LMW thiols have not yet been identified in *Cba. tepidum* or other *Chlorobiaceae*. Prior studies suggested the existence of a novel thiol, named U11, in *Chlorobium limicola*, which also lacked detectable amounts of GSH and other common LMW thiols (18).

Here we show that *Cba. tepidum* contains N-methyl-bacillithiol (N-Me-BSH) consisting of BSH modified by N-methylation of the cysteine motif. Orthologs of BSH biosynthesis genes present in the *Cba. tepidum* genome are required for the synthesis of N-Me-BSH as is a SAM methyltransferase encoded by CT1040 that performs the methylation of BSH. N-Me-BSH was detected in all members of the *Chlorobi* examined. Orthologs of BSH biosynthesis genes and CT1040 co-occur in the genomes of extremely diverse Bacteria, and the presence or absence of a CT1040 ortholog was shown to accurately predict the thiol content in *Polaribacter* sp. strain MED152 and *Thermus thermophilus* HB27. The distribution of these genes suggests that BSH and/or N-Me-BSH may be the most widespread LMW thiols in biology.

## RESULTS

### *Cba. tepidum* contains a novel LMW thiol

LMW thiol compounds in *Cba. tepidum* were examined by HPLC analysis of S-bimane derivatives produced by simultaneous thiol extraction and treatment with the thiol-selective fluorophore monobromobimane (mBBr, Fig. S1). *Cba. tepidum* cells, grown with sulfide plus thiosulfate as electron donors to late-log phase (approx. 20 μg Bchl *c* ml^-1^), produced one bimane derivative with a unique retention time (Fig. 2A arrow) relative to standard thiol compounds and reagent blanks (Fig. S2). The bimane derivative, named U7 for its retention time, was not observed in extracts treated with the sulfhydryl blocking agent N-ethylmaleimide (NEM) prior to mBBr derivatization (Fig. 2B, dashed trace). The observed retention time for this compound was unique relative to all authentic standards analyzed here and to those reported in the literature for other LMW thiol compounds (1, 3, 17–20). For example, extraction of *Escherichia coli*, where glutathione is the predominant LMW thiol (21), yielded a mBBr derivative that co-migrated with a glutathione standard and was absent from *Cba. tepidum* extracts (Fig. 2C). This result confirms earlier reports that other members of the *Chlorobi* lack GSH (18).

**Figure 2.**
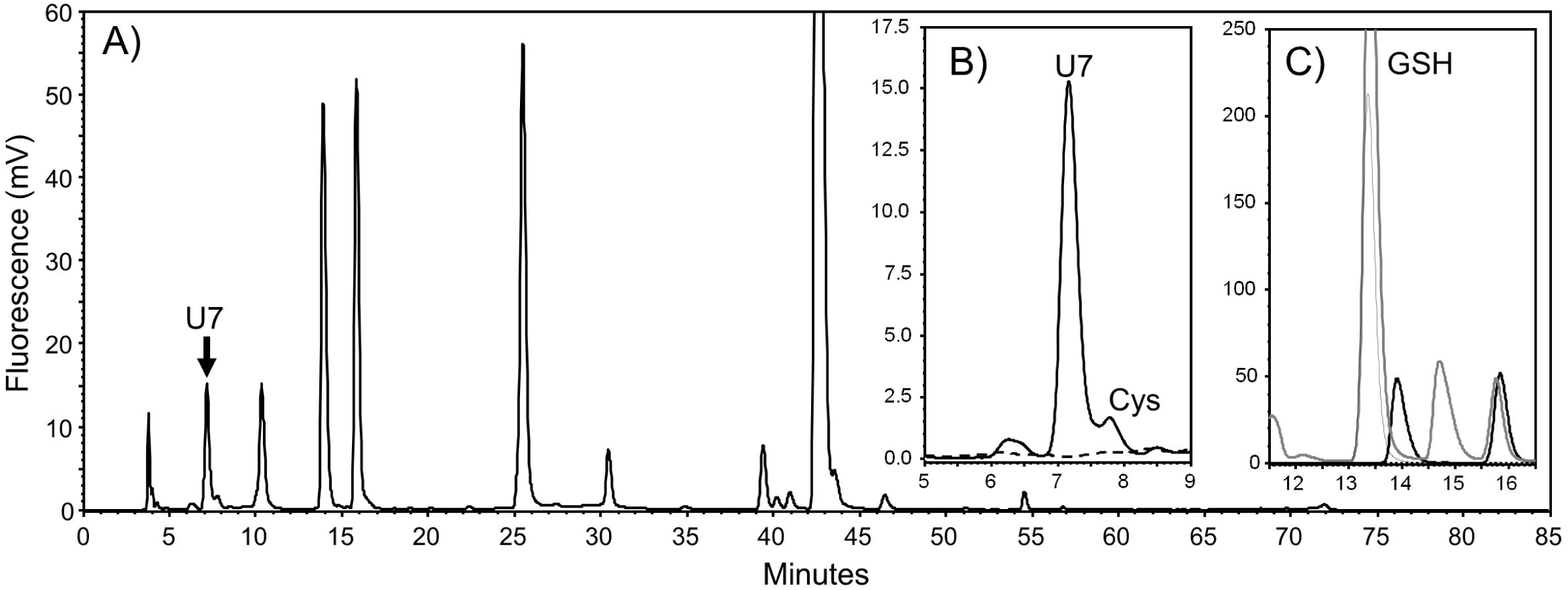
*Cba. tepidum* contains a novel LMW thiol. (A) Full HPLC chromatogram of bimane extract from a stationary phase *Cba. tepidum* culture grown under standard conditions. The arrow indicates the novel thiol U7. (B) U7 and cysteine (Cys) are not detected if extract is treated with N-ethyl-maleimide (dashed line) before mBBr. (C) Glutathione (purified standard, dash-dotted line) is readily detected in *E. coli* bimane extract (grey line), but not *Cba. tepidum* (solid line).

### U7 is BSH modified by N-methylation of cysteine

Treating bimane-labelled extracts with TCEP according to Franz *et al.* (22) did not change the U7 peak area (Fig. S3) demonstrating that U7 is a monothiol (R-SH) rather than a perthiol (R-S-S_n_-H). HPLC fractions containing U7 were collected, concentrated, and analyzed by Fourier transform ion cyclotron resonance mass spectrometry (FT-ICR-MS) in positive ion mode yielding a predominant ion at 603.196 *m/z* for two independent samples (Fig. S4, Table S1). This is consistent with a mass of 412 Da for the thiol before addition of the bimane tag (191 Da). DECOMP analysis of the FT-ICR-MS data for monoisotopic element combinations within 2 ppm error and formulas containing at minimum: bimane (C_10_H_11_N_2_O_2_), one sulfur atom, and 10 more carbon atoms produced 13 possible formulas. The composition of bimane was replaced with hydrogen to arrive at likely formulas for the original thiol (Table S2, first column). Candidates were further evaluated by comparison to MS/MS data. MS^3^ data on the 469 *m/z* ion indicated that masses observed in MS^2^ of the 603 *m/z* ion were produced by sequential decomposition events: 603>469>433>391 *m/z.* Only one formula, C_14_H_24_N_2_O_10_S, could produce the correct decomposition masses given the starting formula (Table S2, bold text): U7 > *a*-C_10_H_18_N_2_O_5_S > *b*-C_10_H_14_N_2_O_3_S > *c*-C_8_H_12_N_2_O_2_S.

Fragment *a* indicated a loss of malic acid (C_4_H_6_O_5_) from the U7 bimane adduct. Malic acid addition to UDP-N-acetyl-glucosamine (UDP-Glc-NAc) is the first step of BSH biosynthesis (3), suggesting that U7 could be related to BSH.

The deduced mass and formula for U7 differs from BSH by an additional methylene unit. Two plausible structures that could account for this are a BSH derivative where the cysteine sidechain is replaced with either homocysteine (hCys-BSH) or N-methylcysteine (N-Me-BSH). To address this, S-bimane derivatives of hCys-BSH and N-Me-BSH were chemically synthesized as analytical reference samples for comparison with the bimane labelled U7 by HPLC separation (see Supporting Information). U7mB extracted and purified from *Cba. tepidum* co-migrated with N-Me-BSmB, but not hCys-BSmB, when analyzed in separate runs and a single symmetrical peak when U7mB was spiked with synthetic N-Me-BSmB (Fig. 3). Therefore, we conclude that *in vivo* U7 is N-Me-BSH. N-Me-BSH could be quantified in *Cba. tepidum* and other *Chlorobiaceae* (Table 1): *Chlorobium phaeobacteroides* DSM265 and *Prosthecochloris* sp. strain CB11, which was recently enriched from the Chesapeake Bay (28). N-Me-BSH was also observed in *Chlorobium luteolum* DSM273 and *Prosthecochloris aestuarii* DSM271 at ~2-8 pmol (mg dw)^-1^.

**Table 1.** Detection of N-Me-BSH and BSH in selected bacteria grown to early stationary phase.

|  |  | *nmbA* | [N-Me-BSH] | [BSH] |
| --- | --- | --- | --- | --- |
| Organism |  | ortholog | pmol (mg dw)<sup>-1</sup> | pmol (mg dw)<sup>-1</sup> |
| *Cba. tepidum* | wild type | CT1040 | 592±373 | ~200<sup>a</sup> |
| “ | ΔCT1040 (*nmbA*) | none | B.D.L. | 791±157 |
| *Prosthecochloris* sp. | strain CB11 | ?<sup>c</sup> | 65±29 | B.D.L. |
| *Chlorobium phaeovibrioides* | DSM265 | Cvib_0902 | 109±10 | B.D.L. |
| *Polaribacter* sp. | MED152 | MED152_02425 | 1,151±392 | ~200<sup>a</sup> |
| *Thermus thermophilus* | HB27 | N.D.<sup>d</sup> | B.D.L. | 27±9 |
| 518 | a-Quantification of BSH at low levels is inaccurate due to an earlier eluting, overlapping |  |  |  |
| 519 | peak in cell extracts from these strains. |  |  |  |
| 520 | b-Below detection limit, ~0.5 pmol (mg dw)<sup>-1</sup> for N-Me-BSH and 20 pmol (mg dw)<sup>-1</sup> for |  |  |  |
| 521 | BSH |  |  |  |
| 522 | c-No genome sequence is available for this organism |  |  |  |
| 523 | d-No bidirectional BLASTP best-hit with e-value < 1e-30 was detected |  |  |  |

**Figure 3.**
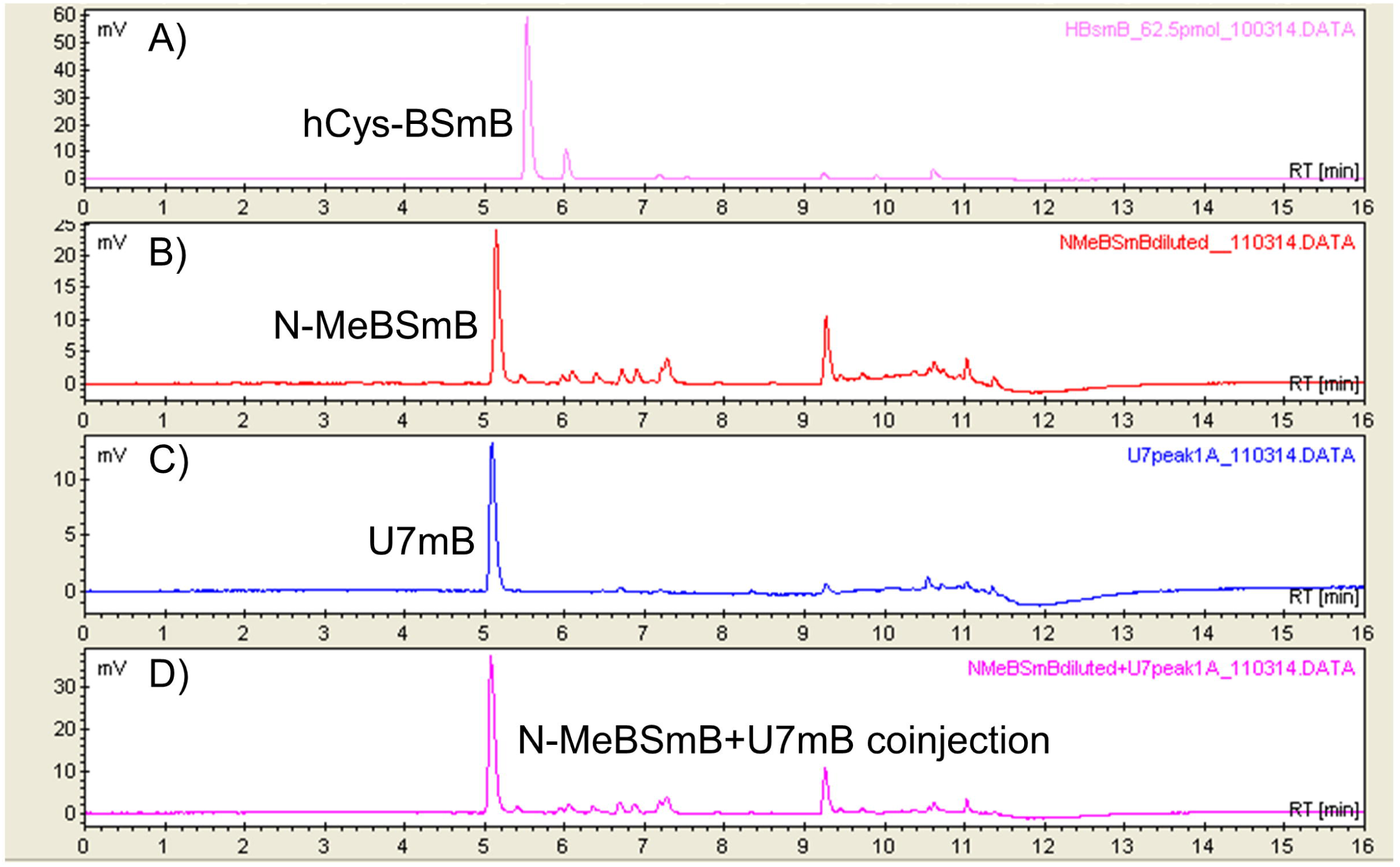
The novel *Cba. tepidum* U7-bimane adduct (U7mB) is N-Me-BSmB. HPLC chromatograms of: (A) synthetic hCys-BSmB, (B) synthetic N-Me-BSmB, (C) as purified *Cba. tepidum* U7mB, (D) mixture of N-Me-BSmB and as purified *Cba. tepidum* U7mB. The retention times are different from Fig. 1 because a different column and elution gradient were used to better separate these compounds.

### Genetic identification of the *Cba. tepidum* N-Me-BSH biosynthetic pathway

All *Chlorobiaceae* all genome sequences encode orthologs of the three enzymes required for BSH biosynthesis from UDP-Glc-NAc, malic acid and cysteine in *Bacillus subtilis* (Fig. S5). BshA (CT0548 in *Cba. tepidum*) condenses UDP-Glc-NAc and malic acid to produce D-Glc-NAc-L-Mal that is hydrolyzed to D-GlcN-L-Mal by BshB (CT1419), which is condensed with cysteine by BshC (CT1558) to produce BSH (Fig. 4A) (23). The requirement for this pathway for N-Me-BSH synthesis was confirmed by deleting CT1419/*bshB* from the *Cba. tepidum* genome. The resulting mutant strain did not contain detectable levels of N-Me-BSH (Fig. 4B, Table 1) or BSH (Table 1).

**Figure 4.**
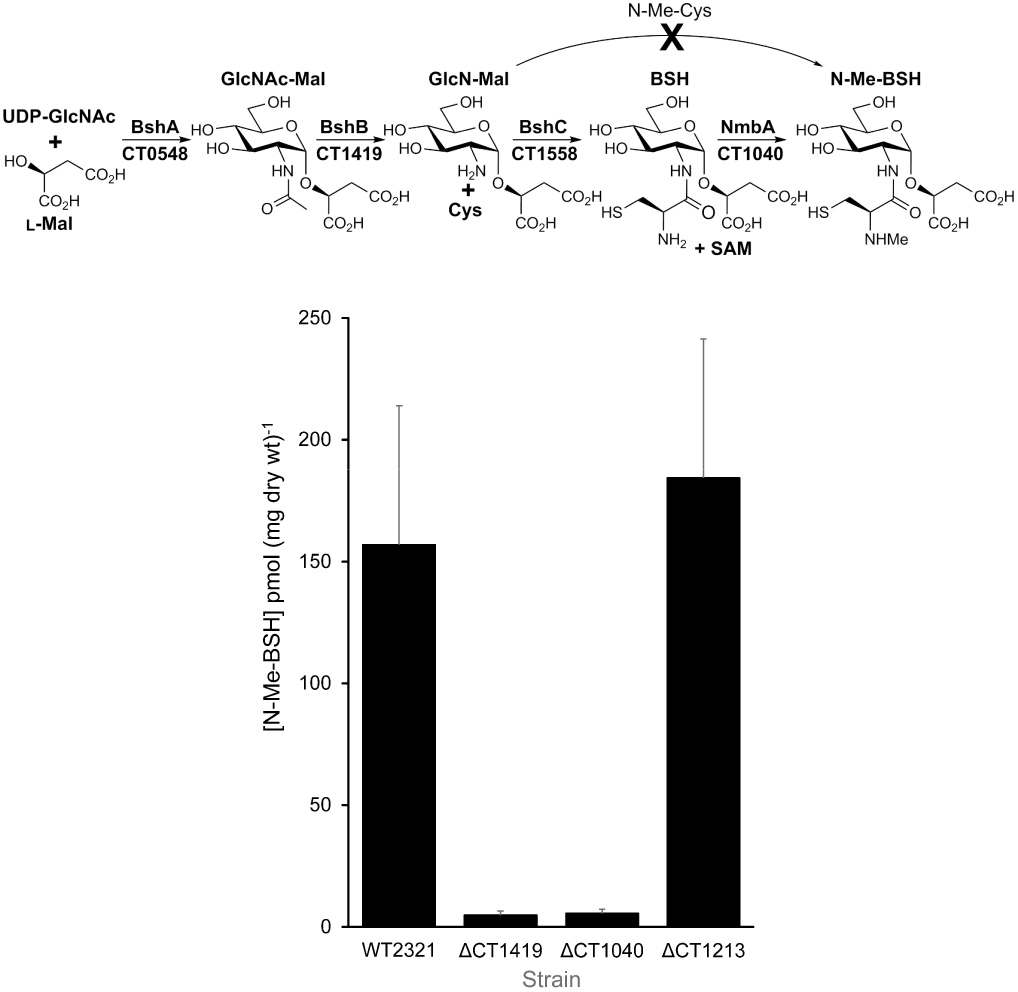
The proposed biosynthetic pathway for N-Me-BSH (A) and N-Me-BSH pool size (B) in *Cba. tepidum* deletion mutant strains ΔCT1419 (*bshB*), ΔCT1040 (putative SAM-dependent methyltransferase) and ΔCT1213 (putative SAM-dependent methyltransferase). The values shown for strains ΔCT1419 and ΔCT1040 indicate the limit of detection for N-Me-BSH. There is currently no evidence to support N-Me-BSH synthesis via N-Me-Cys addition to GlcNMal.

N-Me-BSH biosynthesis could potentially proceed via the ligation of N-Me-cysteine with GlcN-Mal or the N-methylation of BSH. BSH is often co-detected with N-Me-BSH in *Cba. tepidum* extracts (Table 1), but N-Me-cysteine has never been observed (data not shown). This suggests a BSH methyltransferase is required to synthesize N-Me-BSH. Orthologs of two putative SAM-dependent methyltransferases in the *Cba. tepidum* genome, CT1040 and CT1213, are found in all *Chlorobi* genomes as are genes encoding BSH biosynthesis (Fig. S5). Each gene was deleted from the *Cba. tepidum* genome and the resulting mutant strains analyzed for LMW thiols. The strain lacking CT1040 did not contain N-Me-BSH while the strain lacking CT1213 contained similar levels of N-Me-BSH as the wild type (Fig. 4B). Furthermore, the strain lacking CT1040 contained levels of BSH similar to the concentration of N-Me-BSH in the parental wild type strain (Table 1) suggesting that the deletion of CT1040 causes a complete blockage of BSH methylation in this strain. Therefore, we conclude that the CT1040 gene product functions *in vivo* as a SAM-dependent BSH methyltransferase for which we propose the name NmbA for <u>N</u>-<u>M</u>e-<u>B</u>SH synthase <u>A</u>.

### N-methyl-BSH pool size varies with physiological status and is in the reduced state in vivo

*Cba. tepidum* was grown with a variety of sulfur compounds as electron donors for photosynthesis and the N-Me-BSH pool size quantified at different growth stages. N-Me-BSH pool sizes increased during growth and were always highest in stationary phase (45 hrs, Fig. 5A). N-Me-BSH pool size and biomass were strongly correlated regardless of the electron donor used for growth (r^2^ = 0.788). *Cba. tepidum* cultures grown with thiosulfate and sulfide had the highest biomass concentrations followed by thiosulfate and sulfide. This suggests that as cell density increases, the decrease in light intensity due to self-shading may drive an increase in the N-Me-BSH pool size. This was supported by the fact that *Cba. tepidum* grown at low light intensity contained five-fold more N-Me-BSH compared to cells grown at standard or high light intensity (Fig. 5B).

**Figure 5.**
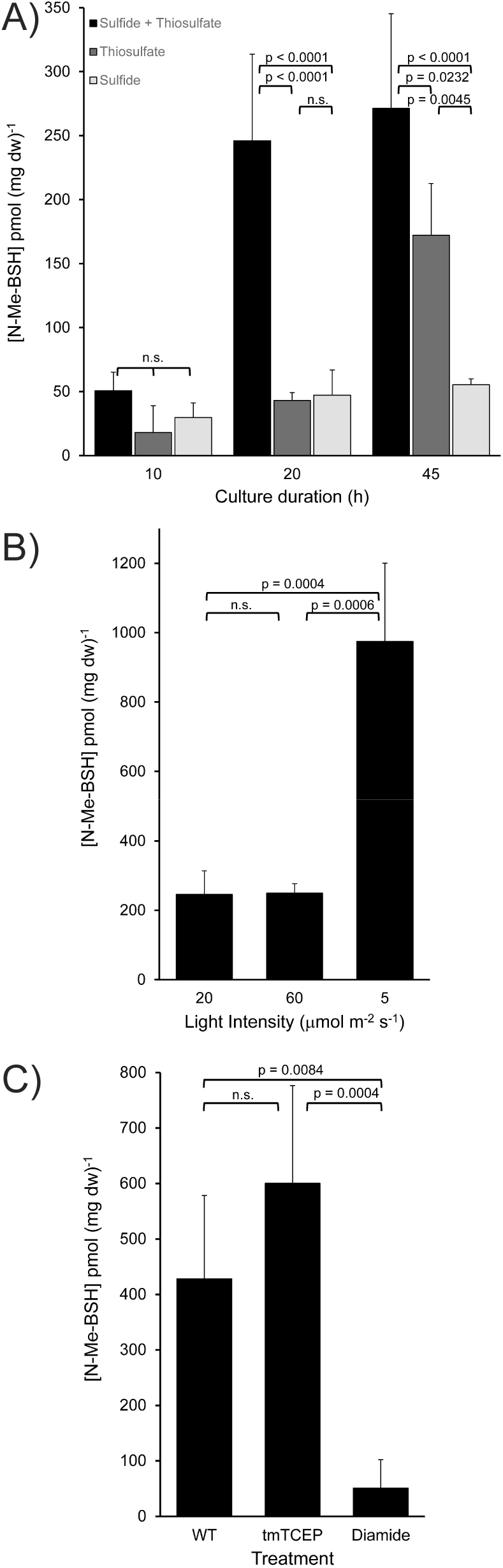
Dynamics of N-Me-BSH pool sizes in *Cba. tepidum.* (A) Pool size in *Cba. tepidum* grown for the indicated times (mid log phase, late log phase, early stationary phase) with different electron donor combinations. (B) Pool size in the wild type grown with the indicated light fluxes grown to stationary phase. (C) Pool size in stationary phase wild-type cells or cells treated with 1 mM trimethyl-TCEP (tmTCEP) or 2 mM diamide. Significant differences are indicated by p-values calculated by the Tukey-Kramer HSD test after ANOVA where “n.s.” indicates not significant (p > 0.05).

The redox state of the N-Me-BSH pool was assessed by treating stationary phase *Cba. tepidum* cultures with trimethyl-TCEP, a phosphine reductant that is able to cross phospholipid bilayers (24), or diamide, a disulfide-generating electrophile that is used to induce sulfhydryl specific oxidative stress (25–27). Addition of trimethyl-TCEP increased the N-Me-BSH pool size 1.4-fold compared to compared to the untreated culture, but this change was not significant (Fig. 5C). In contrast, the addition of diamide decreased the N-Me-BSH pool size 8.4-fold (Fig. 5C). Together these results demonstrate that N-Me-BSH is found predominantly in its reduced state in cells, and is a redox responsive thiol in *Cba. tepidum.*

### Phylogenetic distribution of LMW thiol biosynthetic genes

The direct detection and structural analysis of LMW thiol metabolites is the gold standard for assessing their distribution (1, 28). The current distribution of directly detected LMW thiols in bacteria is outlined in Figure 6A. The *Polaribacter* sp. strain MED152 (*Bacteroidetes*) genome contains orthologs of BSH biosynthetic genes *bshA*-*C* and CT1040/*nmbA*, while the *Thermus thermophilus* HB27 (*Deinococcus*-*Thermus*) genome only contains *bshA*-*C*. HPLC analysis of bimane labeled cell extracts clearly showed that *Polaribacter* sp. strain MED152 contained high levels of N-Me-BSH while *T. thermophilus* HB27 contained BSH, but not N-Me-BSH (Table 1). Based on this clear relationship between gene and LMW thiol content, all complete microbial genome sequences in the Integrated Microbial Genomes database were searched for the presence of orthologs of *bshA*-*C* and *nmbA.* A complete N-Me-BSH biosynthetic pathway is found not only in the *Chlorobi* and *Bacteroidetes*, but also in members of the phyla *Acidobacteria*, *Firmicutes*, and in a basal member of the *Chlamydiae*, *Waddlia chondrophila* (Fig. 6B). BSH biosynthesis has been chemically demonstrated in the *Deinococcus*-*Thermus* lineage and *Firmicutes* (3), this analysis predicts BSH would also be found in members of the *Bacteroidetes* and *Acidobacteria.* In comparison, mycothiol biosynthesis genes (*mshA*-*D*) are only found in the *Actinobacteria* and glutathione biosynthesis genes (*gshA*-*B*) in one member of the *Actinobacteria* (*Frankia* sp. EAN1pec), the *Cyanobacteria*, *Proteobacteria*, and *Eukarya.*

**Figure 6.**
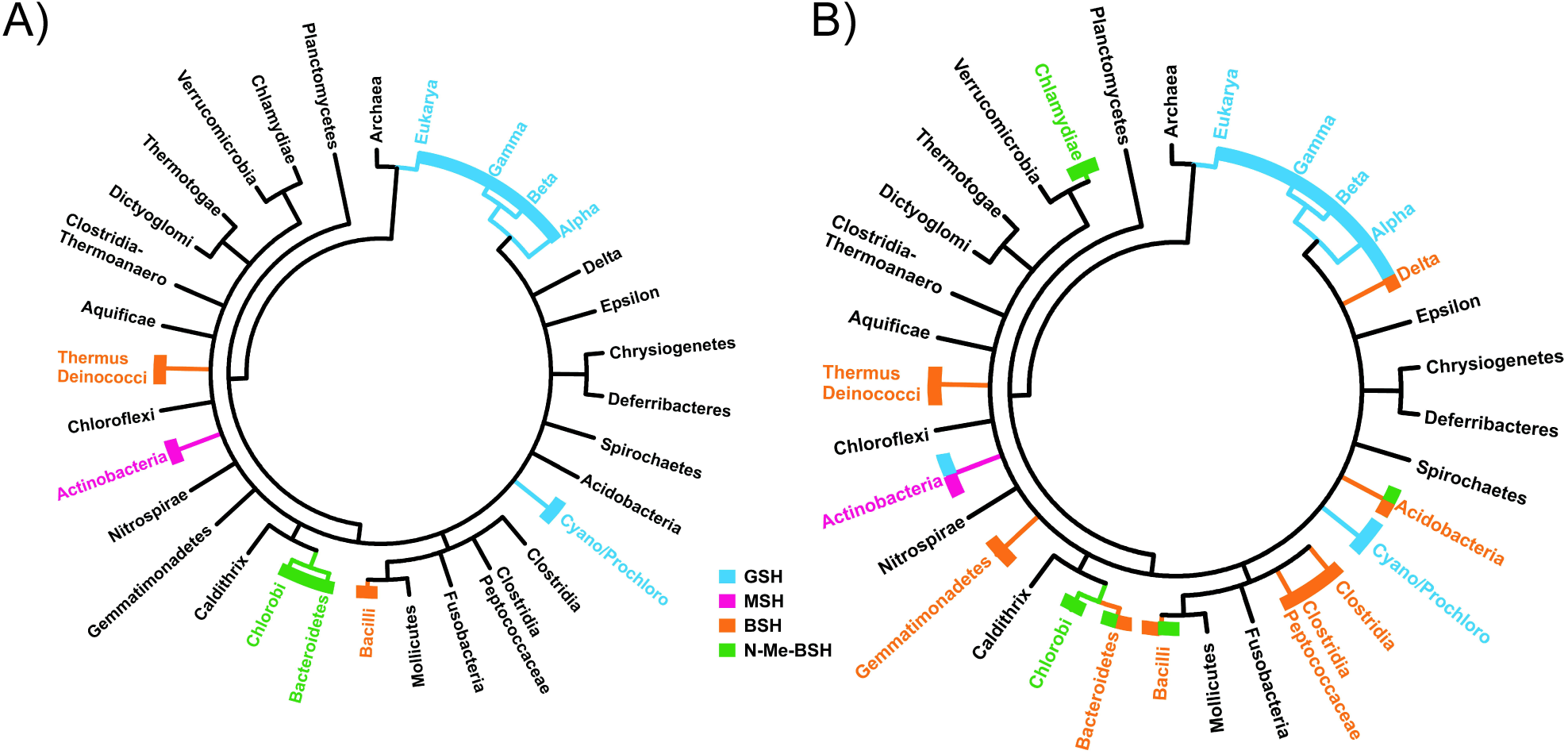
The distribution of LMW thiols in bacteria as determined by mBBr derivatization-HPLC (A) and the potential distribution based on an analysis of complete genome sequences (B) for the presence orthologs encoding complete pathways for GSH (*gshA*-*B*), MSH (*mshA*-*D*), BSH (*bshA*-*C*) or N-Me-BSH (*bshA*-*C* + *nmbA*).

Within the *Chlorobi* and sister phylum *Ignavibacteriae*, all genomes appear to have a complete N-Me-BSH pathway except for the draft genome of NICIL-2, likely the most basal member of the *Chlorobi* (29), that contains *bshA*-*C*, but not CT1040/*nmbA*. Outside of these taxa, around a third of the BSH-pathway positive *Acidobacteria* (33%) and *Bacteroidetes* (31%) contain an *nmbA* ortholog while smaller numbers of *Firmicutes* appear to possess *nmbA* (7%, multiple *Paenibacillus* spp. and *Brevibacillus brevis*).

## DISCUSSION

The data presented here show that N-Me-BSH is the major LMW thiol in the *Chlorobiaceae*. N-Me-BSH biosynthesis requires the function of gene products orthologous to *bshA*-*C* and a SAM-dependent methyltransferase named *nmbA* with CT1040 as the defining member of this gene family. The N-methylation of cysteine in secondary metabolites is rarely observed in biology. To the best of our knowledge, N-Me-BSH is only the fourth example of this modification. The others are pyochelin, an N-methylated thiazolidine containing siderophore produced by *Pseudomonas aeruginosa* (30), kendarimide A, a poly N-methylated oligopeptide including N-Me-Cys isolated from the Indonesian sponge *Haliclona* sp. (31), and thiocoraline; a depsipeptide isolated from a marine *Micromonospora* sp. (32). Pyochelin is, and the others are likely to be, synthesized by extremely large multi-domain non-ribosomal peptide synthetases where a specific MTase domain catalyzes the cysteine modification (30). N-Me-BSH appears to be the first case of cysteine N-methylation outside of oligopeptide metabolites where the N-methylation is catalyzed by a standalone MTase.

The functional consequences of producing N-Me-BSH vs. BSH are currently unknown. While BSH is often detected in N-Me-BSH producing bacteria, N-Me-BSH levels are always greater than those of BSH. The concentrations of N-Me-BSH detected in the *Chlorobi* under normal anaerobic growth conditions, 65-700 pmol thiol (mg dw)^-1^, are at the lower end of the range of the BSH values previously reported in various *Bacilli* and *Deinococcus radiodurans*, 200-2,600 pmol thiol (mg dw)^-1^ (3, 33). The elevation in N-Me-BSH levels during exponential growth and reaching a maximum in stationary phase is similar to observations of BSH in *B. subtilis* (34). However, low light growth elevated the N-Me-BSH level to ~1,050 pmol thiol (mg dw)^-1^ in *Cba. tepidum*, similar to the level of N-Me-BSH in aerobically grown *Polaribacter* sp. strain MED152, ~1,150 pmol (mg dw)^-1^. Environmental conditions clearly influence intracellular levels of N-Me-BSH, which will warrant further investigation.

Modification of their predominant LMW thiol structure during stationary phase is known in some bacteria. Some marine actinomycetes produce MSH with a N-propionyl group instead of the normal N-acetyl group. The diversion of propionyl-CoA into N-propionyl-MSH is proposed to limit propionyl-CoA accumulation during degradation of odd chain and branched chain fatty acids (35). *E. coli* have also been shown to convert much of their GSH pool to glutathionylspermidine during stationary phase under anaerobic conditions, which is believed to modulate free GSH and/or spermidine in response to different environmental conditions (36). However, the *Chlorobiaceae* predominantly make N-Me-BSH irrespective of growth phase suggesting that there has been a selection for this molecule in their physiology.

Physiologically, N-Me-BSH is the best current candidate for a LMW thiol proposed to facilitate the trafficking of sulfur atoms between the periplasm and cytoplasm in phototrophic sulfur oxidizing bacteria, a role proposed for glutathione amide in *C. gracile* (14). However, the fact that *Polaribacter* sp. MED152, which does not employ sulfur-based energy metabolism, synthesizes N-Me-BSH and many other members of the *Bacteroidetes*, *Acidobacteria* and *Firmicutes* carry *nmbA* orthologs and likely synthesize N-Me-BSH indicate that the molecule cannot be exclusively tied to sulfur oxidation. Furthermore, in dissimilatory sulfate reduction, the DsrC protein has been shown to stimulate the activity of dissimilatory sulfite reductase, DsrAB, and appears to act as the preferred acceptor for the reduced sulfur atom by forming a trisulfide bridge between two cysteine side chains (37). DsrC-trisulfide is then proposed to be reduced by DsrMKJOP generating sulfide and regenerating DsrC to accept another sulfur atom. As phototrophic sulfur oxidation is proposed to involve a reversed Dsr system for oxidizing elemental sulfur to sulfite (15, 16), sulfur atom transfer to DsrC as a cytoplasmic acceptor may operate in place of a LMW thiol shuttle. This would explain the lack of correlation between N-Me-BSH and sulfur-based energy metabolism in these organisms.

Another possible function in the *Chlorobiaceae* is that N-Me-BSH is the *in vivo* reductant for the recently described excitation energy regulating mechanism in the FmoA protein (38). Cysteine centered thiyl radicals in FmoA are proposed to interact with excited bacteriochlorophyll *<u>a</u>* to prevent excitation energy transfer to the reaction center under unfavorable conditions. *In vitro*, dithionite, dithiothreitol, sulfide, glutathione and TCEP were capable of capable of regulating FmoA energy transfer, and GSH was proposed as the *in vivo* mediator (38). The results presented here indicate that GSH is not a good candidate as no *Chlorobiaceae* genomes encode GSH biosynthesis (Fig. 6B, Dataset S1) and GSH was not detected in *Cba. tepidum* (Fig. 1), *Chl. phaeobacteroides* DSM265, *Prosthecochloris* sp. strain CB11, *Chl. luteolum* DSM273, and *P. aestuarii* DSM271. Rather, these data suggest N-Me-BSH as the most likely thiol based redox modulator of FmoA energy transfer. The data here predict that *Chloracidobacterium thermophilum* (39), the only organism outside of the *Chlorobi* to utilize FmoA in light harvesting, should synthesize N-Me-BSH (Dataset S1) bolstering this assertion. BSH based thiols may be more suitable for this function because they have significantly lower redox potential than GSH (34). Detailed examinations of the physical and redox properties of N-Me-BSH and the mutant strains generated here will help to address this question. However, as with sulfur-based energy metabolism, the occurrence of N-Me-BSH in organisms that do not contain FmoA means that N-Me-BSH cannot be exclusively associated with light harvesting.

The genetic data indicate that N-Me-BSH is synthesized in *Cba. tepidum* after BSH biosynthesis by the CT1040/*nmbA* gene product, a SAM-dependent methyltransferase. The role of methylation can thus be explored by generating mutant strains expressing non-cognate thiols, i.e. N-Me-BSH in *B. subtilis* and BSH in *Cba. tepidum*, to address the functional significance of this rare metabolic modification. As a standalone methyltransferase, NmbA could potentially be used to methylate a wide range of small molecule targets to improve properties or activities. Identifying *nmbA* allowed us to predict and confirm LMW thiol biosynthetic capacity in complete genome sequences over long phylogenetic distances. This, in turn, led us to conclude that LMW thiols based on the BSH backbone are likely the most widely distributed thiols in biology. The analysis also highlighted groups of organisms that should be targeted to more fully understand the diversity of LMW thiol structure and function. Major bacterial lineages e.g. *Verrucomicrobia*, *Planctomycetes*, *Spirochates* and others, have no documented or predicted LMW thiol for redox homeostasis, a critical cellular function. Future genome directed studies of LMW thiol diversity, structure and function may uncover further variations on LMW thiol molecular backbones that underlie critical metabolic processes and where enzymes generating this biochemical diversity may find applications for engineered product synthesis.

## MATERIALS AND METHODS

### Bacterial growth conditions and media

All strains and antibiotic selections used in this study are listed in Table S3. *Escherichia coli* strains were grown in Lysogeny Broth at 37 °C (40). *Chlorobaculum tepidum* strains were grown in Pf-7 medium buffered to pH 6.95 with the addition of 1,3-bis(tris(hydroxymethyl)methylamino)propane (BTP, MP Biomedicals, Solon, OH) as previously described (7, 41) in 250 ml narrow neck screw cap media bottles sealed with black open top phenolic caps containing a flanged butyl rubber septum (Fisher Scientific, Pittsburgh, PA). All cultures were maintained anaerobically and pressurized to 10 psi with 5% CO_2_/95% N_2_. *Cba. tepidum* cultures were grown at 47 °C with 20 μmol photons m^-2^ s^-1^ (standard light), 60 μmol photons m^-2^ s^-1^ (high light), or 5 μmol photons m^-2^ s^-1^ (low light) of irradiance, supplied by 40 or 100 W neodymium full-spectrum bulbs (Lumiram Electric Corp., Larchmont, NY). All irradiance measurements were made with a light meter equipped with a quantum PAR sensor (LI-COR, Lincoln, NE).

### Metabolite extraction and mBBr derivatization

A modified version of the mBBr extraction and derivatization protocol of Fahey and Newton (19) was used to extract thiols from both *Cba. tepidum* and *E. coli* (Fig. S1). Details on equipment and HPLC separations are in provided in Supporting Information.

### Effect of reductant or oxidant on LMW thiol pool size

Stationary phase wild-type *Cba. tepidum* cells (48 hrs of growth) were treated with 2 mM trimethyl-TCEP, synthesized as described in Supporting Information (24), 1 mM diamide, or nothing to cultures that were then incubated for 40 min in an anaerobic chamber before samples were harvested for mBBr extraction and derivatization.

### Liquid chromatography-tandem mass spectrometry

Bimane derivatives of interest were collected from multiple HPLC runs of the same sample and lyophilized (Labconco FreeZone 4.5, Kansas City, MO) for 12 hours. The concentrated material was resuspended in 0.075% (v/v) glacial acetic acid and 68% (v/v) methanol followed by further concentration in a SpeedVac and reconstitution in water.

High resolution, positive ion mode ESI LC-FTICR-MS was performed with a 50 mm C18 column. Solvent A was 0.1% aqueous acetic acid, pH 3.5, and Solvent B was methanol. The 30 min elution protocol (0.2 ml min^-1^) was as follows: 0 min, 15% B; 5 min, 15% B; 15 min, 23% B; 17 min, 42% B; 20 min, 42% B; 20.02, 15% B. This method and solvent system resulted in one bimane derivative eluting at 5.7 min which was then analyzed by a 7 T Fourier-transform ion cyclotron resonance mass spectrometer (FT-ICR MS) from Thermo Scientific (LTQ FT Ultra hybrid mass spectrometer). The front-end of the instrument is a linear ion trap mass spectrometer (LTQ MS) which serves as the ion accumulation site for ultra-high resolution analysis in the ICR cell. Fragmentation was provided by collision-induced dissociation (CID) in the linear ion trap followed by FT-ICR MS analysis.

### Synthesis and analysis of bacillithiol derivatives

S-bimane derivatives of N-Me-BSH and hCys-BSH were synthesized following similar procedures previously developed for BSH synthesis (42), but using suitably protected N-methyl cysteine and homocysteine building blocks in place of the protected cysteine building block used for BSH synthesis (see Supporting Information).

### Gene deletion and mutant analysis

Full details of the protocol for in-frame deletion of genes in *Cba. tepidum* will be described elsewhere (Hilzinger, Raman and Hanson, in preparation). Briefly, regions flanking the gene to be deleted were PCR amplified using primers listed in Table S4 and inserted into pKO2.0-Sm/Sp by Gibson Assembly (New England Biolabs, Ipswich, MA). Plasmid pKO2.0-Sm/Sp is based on pKO2.0 used to delete genes in *Shewanella oneidensis* (43). pKO2.0 was modified by replacing the gentamycin resistance gene with the streptomycin/spectinomycin resistance cassette from pHP45Ω (44). Plasmids were mobilized from *E. coli* strain β-2155 to *Cba. tepidum* by conjugation (12, 43). Single recombinants were selected with spectinomycin and streptomycin, verified to contain both the WT and deletion alleles of the gene by PCR and then grown in Pf-7 medium without antibiotic selection. Double recombinants were obtained by plating this culture onto solid medium containing 10% w/v sucrose and strains containing only the deletion allele of the gene of interest were identified by PCR. These were grown in liquid Pf-7 medium for thiol analysis as described above.

### Bioinformatic analysis of LMW thiol biosynthetic pathways

Orthologs of genes for LMW thiol biosynthetic pathways were collected from finished microbial genome sequences in the Integrated Microbial Genomes database (https://img.jgi.doe.gov) using the Custom Homolog Selection tool requiring a minimum amino acid sequence identity of 30%, a minimum BLASTP e-value of 1^-20^, and similar length of query and subject. Genes used as queries to collect the orthologs were CT0548 (*bshA*), CT1419 (*bshB*), CT1558 (*bshC*), b2688 (*gshA*), b2947 (*gshB*), slr0990 (*gshA*), slr1238 (*gshB*), Rv0486 (*mshA*), Rv1170 (*mshB*), Rv2130c (*mshC*), Rv0819 (*mshD*). Taxonomic information was retrieved for each genome containing an ortholog of a given gene and combined lists for genomes containing multiple orthologs (i.e. all genomes containing *mshA* + *mshB* + *mshC* + *mshD*) constituting a pathway assembled from single ortholog lists using the UNIX command “grep” and the LOOKUP function in Microsoft Excel. The complete list of genomes inferred to encode the biosynthesis of each thiol is provided as Dataset S1 in the Supporting Information. SAM-dependent methyltransferases were identified in the *Cba. tepidum* genome by searching for proteins containing domain cd02440 (AdoMet_MTases), but lacking obvious functional annotation. These protein sequences were used as queries for BLASTP searches against all other *Chlorobi* proteins to determine their distribution relative to the *bshA*-*C* orthologs above.

## Acknowledgments

This work was supported by a CAREER award MCB-0447649 from the National Science Foundation to T.E.H, a Leverhulme Trust Research Grant RPG-2012-506 to SVS, and utilized common instrumentation facilities provided in part by grant P20-RR116472-04 from the IDeA Networks of Biomedical Research Excellence program of the National Center for Research Resources, National Institutes of Health. The authors would like to thank Yong Seok Choi and Kelvin H. Lee for initial LC-MS/MS analysis of purified N-Me-BSH and the Fourier Transform Mass Spectrometry Facility at the Woods Hole Oceanographic Institution for providing the data presented here.

## Author Contributions

JH, TEH, and CH designed research; JH, VR, SVS, RAJT, MA, DFR, JNM, CH and TEH performed research and analyzed data; JH, RAJT, TEH and CH wrote the manuscript.

## REFERENCES

1. Fahey RC (2013) Glutathione analogs in prokaryotes. Biochim Biophys Acta 1830:3182–3198.

2. Newton GL, Buchmeier N, Fahey RC (2008) Biosynthesis and functions of mycothiol, the unique protective thiol of Actinobacteria. Microbiol Mol Biol Rev 72:471–494.

3. Newton GL, Rawat M, La Clair JJ, Jothivasan VK, Budiarto T, Hamilton CJ, Claiborne A, Helmann JD, Fahey RC (2009) Bacillithiol is an antioxidant thiol produced in Bacilli. Nat Chem Biol 5:625–627.

4. Evans MC, Buchanan BB, Arnon DI (1966) A new ferredoxin-dependent carbon reduction cycle in a photosynthetic bacterium. Proc Natl Acad Sci U S A 55:928–934.

5. Orf GS, Blankenship RE (2013) Chlorosome antenna complexes from green photosynthetic bacteria. Photosynth Res 116:315–331.

6. Oostergetel GT, van Amerongen H, Boekema EJ (2010) The chlorosome: a prototype for efficient light harvesting in photosynthesis. Photosynth Res 104:245–255.

7. Wahlund TM, Woese CR, Castenholz RW, Madigan MT (1991) A thermophilic green sulfur bacterium from New Zealand hot springs, *Chlorobium tepidum* sp. nov. Arch Microbiol 156:81–90.

8. Eisen JA, Nelson KE, Paulsen IT, Heidelberg JF, Wu M, Dodson RJ, Deboy R, Gwinn ML, Nelson WC, Haft DH, Hickey EK, Peterson JD, Durkin AS, Kolonay JL, Yang F, Holt I, Umayam LA, Mason T, Brenner M, Shea TP, Parksey D, Nierman WC, Feldblyum T V, Hansen CL, Craven MB, Radune D, Vamathevan J, Khouri H, White O, Gruber TM, Ketchum KA, Venter JC, Tettelin H, Bryant DA, Fraser CM (2002) The complete genome sequence of *Chlorobium tepidum* TLS, a photosynthetic, anaerobic, green-sulfur bacterium. Proc Natl Acad Sci U S A 99:9509–9514.

9. Azai C, Harada J, Hirozo O (2013) Gene expression system in green sulfur bacteria by conjugative plasmid transfer. PLoS One 8:e82345.

10. Frigaard NU, Bryant DA (2001) Chromosomal gene inactivation in the green sulfur bacterium *Chlorobium tepidum* by natural transformation. Appl Environ Microbiol 67:2538–2544.

11. Hanson TE, Tabita FR (2001) A ribulose-1,5-bisphosphate carboxylase/oxygenase (RubisCO)-like protein from *Chlorobium tepidum* that is involved with sulfur metabolism and the response to oxidative stress. Proc Natl Acad Sci U S A 98:4397–4402.

12. Wahlund T, Madigan M (1995) Genetic transfer by conjugation in the thermophilic green sulfur bacterium *Chlorobium tepidum*. J Bacteriol 177:2583–2588.

13. Wahlund TM, Madigan MT (1993) Nitrogen fixation by the thermophilic green sulfur bacterium *Chlorobium tepidum*. J Bacteriol 175:474–478.

14. Seo D, Sakurai H (2002) Purification and characterization of ferredoxin-NAD(P)(+) reductase from the green sulfur bacterium *Chlorobium tepidum*. Biochim Biophys Acta 1597:123–132.

15. Sakurai H, Ogawa T, Shiga M, Inoue K (2010) Inorganic sulfur oxidizing system in green sulfur bacteria. Photosynth Res 104:163–176.

16. Gregersen LH, Bryant DA, Frigaard N-U (2011) Mechanisms and evolution of oxidative sulfur metabolism in green sulfur bacteria. Front Microbiol 2:116.

17. Bartsch RG, Newton GL, Sherrill C, Fahey RC (1996) Glutathione amide and its perthiol in anaerobic sulfur bacteria. J Bacteriol 178:4742–4746.

18. Fahey RC, Buschbacher RM, Newton GL (1987) The evolution of glutathione metabolism in phototrophic microorganisms. J Mol Evol 25:81–88.

19. Fahey RC, Newton GL (1987) Determination of low-molecular-weight thiols using monobromobimane fluorescent labeling and high-performance liquid chromatography. Methods Enzymol 143:85–96.

20. Newton GL, Arnold K, Price MS, Sherrill C, Delcardayre SB, Aharonowitz Y, Cohen G, Davies J, Fahey RC, Davis C (1996) Distribution of thiols in microorganisms: mycothiol is a major thiol in most actinomycetes. J Bacteriol 178:1990–5.

21. Fahey RC, Brown WC, Adams WB, Worsham MB (1978) Occurrence of glutathione in bacteria. J Bacteriol 133:1126–1129.

22. Franz B, Gehrke T, Lichtenberg H, Hormes J, Dahl C, Prange A (2009) Unexpected extracellular and intracellular sulfur species during growth of *Allochromatium vinosum* with reduced sulfur compounds. Microbiology 155:2766–2774.

23. Gaballa A, Newton GL, Antelmann H, Parsonage D, Upton H, Rawat M, Claiborne A, Fahey RC, Helmann JD (2010) Biosynthesis and functions of bacillithiol, a major low-molecular-weight thiol in Bacilli. Proc Natl Acad Sci U S A 107:6482–6486.

24. Cline DJ, Redding SE, Brohawn SG, Psathas JN, Schneider JP, Thorpe C (2004) New water-soluble phosphines as reductants of peptide and protein disulfide bonds: reactivity and membrane permeability. Biochemistry 43:15195–15203.

25. Pöther D-C, Liebeke M, Hochgräfe F, Antelmann H, Becher D, Lalk M, Lindequist U, Borovok I, Cohen G, Aharonowitz Y, Hecker M (2009) Diamide triggers mainly S thiolations in the cytoplasmic proteomes of *Bacillus subtilis* and *Staphylococcus aureus*. J Bacteriol 191:7520–7530.

26. Loi V Van, Rossius M, Antelmann H (2015) Redox regulation by reversible protein S-thiolation in bacteria. Front Microbiol 6:187.

27. Helmann JD (2011) Bacillithiol, a new player in bacterial redox homeostasis. Antioxid Redox Signal 15:123–133.

28. Newton GL, Fahey RC (1995) Determination of biothiols by bromobimane labeling and high-performance liquid chromatography. Methods Enzymol 251:148–66.

29. Hiras J, Wu Y-W, Eichorst SA, Simmons BA, Singer SW (2016) Refining the phylum Chlorobi by resolving the phylogeny and metabolic potential of the representative of a deeply branching, uncultivated lineage. ISME J. 10:833-845.

30. Patel HM, Walsh CT (2001) In vitro reconstitution of the *Pseudomonas aeruginosa* nonribosomal peptide synthesis of pyochelin: characterization of backbone tailoring thiazoline reductase and N-methyltransferase activities. Biochemistry 40:9023–9031.

31. Aoki S, Cao L, Matsui K, Rachmat R, Akiyama S, Kobayashi M (2004) Kendarimide A, a novel peptide reversing P-glycoprotein-mediated multidrug resistance in tumor cells, from a marine sponge of *Haliclona* sp. Tetrahedron 60:7053–7059.

32. Romero F, Espliego F, Baz JP, Quesada TG De, Grávalos D, Calle FD La, Fernández-Puentes JL (1997) Thiocoraline, a new depsipeptide with antitumor activity produced by a marine *Micromonospora*. I. Taxonomy, fermentation, isolation, and biological activities. J Antibiot (Tokyo) 50:734–737.

33. Handtke S, Schroeter R, Jürgen B, Methling K, Schlüter R, Albrecht D, van Hijum SAFT, Bongaerts J, Maurer K-H, Lalk M, Schweder T, Hecker M, Voigt B (2014) *Bacillus pumilus* reveals a remarkably high resistance to hydrogen peroxide provoked oxidative stress. PLoS One 9:e85625.

34. Sharma S V, Arbach M, Roberts AA, Macdonald CJ, Groom M, Hamilton CJ (2013) Biophysical features of bacillithiol, the glutathione surrogate of *Bacillus subtilis* and other firmicutes. Chembiochem 14:2160–2168.

35. Newton GL, Jensen PR, Macmillan JB, Fenical W, Fahey RC (2008) An N-acyl homolog of mycothiol is produced in marine actinomycetes. Arch Microbiol 190:547–557.

36. Smith K, Borges A, Ariyanayagam MR, Fairlamb AH (1995) Glutathionylspermidine metabolism in *Escherichia coli*. Biochem J 312:465–469.

37. Santos AA, Venceslau SS, Grein F, Leavitt WD, Dahl C, Johnston DT, Pereira IAC (2015) A protein trisulfide couples dissimilatory sulfate reduction to energy conservation. Science 350:1541–1545.

38. Orf GS, Saer RG, Niedzwiedzki DM, Zhang H, McIntosh CL, Schultz JW, Mirica LM, Blankenship RE (2016) Evidence for a cysteine-mediated mechanism of excitation energy regulation in a photosynthetic antenna complex. Proc Natl Acad Sci U S A 113:E4486-E4493.

39. Tank M, Bryant DA (2015) *Chloracidobacterium thermophilum* gen. nov., sp. nov.: an anoxygenic microaerophilic chlorophotoheterotrophic acidobacterium. Int J Syst Evol Microbiol 65:1426–1430.

40. Ausubel FM, Brent R, Kingston RE, Moore DD, Seidman JG, Smith JA, Struhl K eds. (2001) Current Protocols in Molecular Biology (John Wiley & Sons, Inc., Hoboken, NJ, USA) doi:10.1002/0471142727.

41. Chan L-KK, Weber TS, Morgan-Kiss RM, Hanson TE (2008) A genomic region required for phototrophic thiosulfate oxidation in the green sulfur bacterium *Chlorobium tepidum* (syn. *Chlorobaculum tepidum*). Microbiology 154:818–829.

42. Sharma S V, Jothivasan VK, Newton GL, Upton H, Wakabayashi JI, Kane MG, Roberts AA, Rawat M, La Clair JJ, Hamilton CJ (2011) Chemical and chemoenzymatic syntheses of bacillithiol: a unique low-molecular-weight thiol amongst low G + C Gram-positive bacteria. Angew Chem Int Ed Engl 50:7101–7104.

43. Burns JL, DiChristina TJ (2009) Anaerobic respiration of elemental sulfur and thiosulfate by *Shewanella oneidensis* MR-1 requires *psrA*, a homolog of the *phsA* gene of *Salmonella enterica* serovar typhimurium LT2. Appl Environ Microbiol 75:5209–5217.

44. Prentki P, Krisch HM (1984) In vitro insertional mutagenesis with a selectable DNA fragment. Gene 29:303–313.

45. Marcucci E, Bayó-Puxan N, Tulla-Puche J, Spengler J, Albericio F (2008) Cysteine-S-trityl a key derivative to prepare N-methyl cysteines. J Comb Chem 10:69–78.

46. Riddles PW, Blakeley RL, Zerner B (1983) Reassessment of Ellman’s reagent. Methods Enzymol 91:49–60.

